# Subsoil arbuscular mycorrhizal fungal communities in arable soil differ from those in topsoil

**DOI:** 10.1101/212597

**Authors:** Moisés A. Sosa-Hernández, Julien Roy, Stefan Hempel, Timo Kautz, Ulrich Köpke, Marie Uksa, Michael Schloter, Tancredi Caruso, Matthias C. Rillig

## Abstract

Arbuscular mycorrhizal fungi are recognized as important drivers of plant health and productivity in agriculture but very often existing knowledge is limited to the topsoil. With growing interest in the role of subsoil in sustainable agriculture, we used high-throughput Illumina sequencing on a set of samples encompassing drilosphere, rhizosphere and bulk soil, in both top- and subsoil. Our results show subsoil AMF communities harbor unique Operational Taxonomic Units (OTUs) and that both soil depths differ in community structure both at the OTU and family level. Our results emphasize the distinctness of subsoil AMF communities and the potential role of subsoil as a biodiversity reservoir.

---

Arbuscular mycorrhizal fungi (AMF) belong to the monophyletic subphylum Glomeromycotina (Spatafora et al., 2016) and form a symbiotic relationship with the roots of most land plants including the majority of agricultural crops (Smith and Read, 2008). This symbiosis is of great relevance for sustainable agriculture due to its ability to increase productivity (Lekberg and Koide, 2005), nutrient uptake (Smith and Smith, 2011), soil aggregation (Leifheit et al., 2014), and plant protection (Veresoglou and Rillig, 2012). In arable fields, the subsoil (i.e. the soil beneath the plough layer) contains more than two thirds of the soil nutrient pool (Kautz et al., 2013). Thus, understanding how AMF communities vary with depth and what factors drive their community assembly is a prerequisite for developing appropriate subsoil management strategies.

Previous studies have shown that AMF spore abundance and diversity in agricultural fields decrease with soil depth (Muleta et al., 2008; Oehl et al., 2005; Yang et al., 2010), however under some circumstances subsoil spore diversity may be greater than in the topsoil (Oehl et al., 2005). Moreover, some species’ spores seem to be associated with particular soil layers, suggesting a vertical variation in community composition (Muleta et al., 2008; Oehl et al., 2005; Yang et al., 2010). These spore-based results have recently been supported by small ribosomal subunit (SSU) cloning-sequencing approaches, finding differences in community composition between AMF communities at different depths (Moll et al., 2016). Nonetheless, we are just starting to unearth subsoil AMF diversity, and the community assembly processes remain unknown.

To shed light on subsoil AMF communities, we used Illumina MiSeq sequencing to analyze a set of soil samples encompassing drilosphere (soil directly influenced by earthworms), rhizosphere (soil directly influenced by roots) and bulk soil (without roots or earthworm burrows), in both top- (10–30 cm, i. e. above the ploughing layer) and subsoil (60–75 cm). Sampling the different compartments was intended to add greater resolution to our results. The first 10 cm were not sampled due to difficulties in differentiating the mentioned compartments at this depth. Samples were collected in May 2011 in a field planted with Cichorium intybus L. (see Uksa et al. (2014) for details).

We performed amplicon-based AMF specific metabarcoding using the AMF specific primer sets described in Krüger et al. (2009) and the primers LR3 and LR2rev (Hofstetter et al., 2002). The final product of amplification is a 350–440 bp region in the LSU including the variable D1-D2 region. Samples were paired-end sequenced on an Illumina MiSeq platform. Bioinformatics details are given in the supplementary material. Briefly, following quality filtering and chimera removal, sequences were de novo clustered at a 97% similarity level into operational taxonomic units (OTUs). Representative sequences of these OTUs have been deposited at ENA under accession numbers LT855246-LT855309.

Taxonomic assignment of the OTUs was carried out using BLAST+ (Camacho et al., 2009) against Glomeromycotina (i.e. AMF) reference sequences published in Krüger et al. (2009) and against the EMBL nucleotide database (Kanz et al., 2005). Following a similar approach as in Martínez-García et al. (2015) for SSU sequences, we considered matches with ≥ 97% similarity a species level match, ≥90% a genus level match, ≥80% a family level match and ≥70% a subphylum level match. A species level match refers to how confidently we assign a name to our OTU based on known sequences, and does not imply that these OTUs are to be considered equivalent to those species. In total, 64 OTUs were confidently assigned to the subphylum Glomeromycotina. Of these, we were able to assign 11 OTUs at the species level, 34 at the genus level, 16 at the family level and 3 at the subphylum level.

The resulting OTU table was analyzed with R version 3.3.1 (R Core Team, 2016). Relative abundance data were obtained by rarefaction of all the samples to the lowest number of reads in a sample (34377 reads), by random subsampling without replacement. Details on the analyses are given in the supplementary material.

Our results present high-throughput molecular evidence that the subsoil AMF community is not simply a subset of the topsoil community, but harbors unique OTUs and that the two soil depths differ in structure both at the family level (Fig. 1) and at the OTU level (Fig. 2). We detected 64 Glomeromycotina OTUs belonging to 7 families (Ambisporaceae, Archaeosporaceae, Claroideoglomeraceae, Diversisporaceae, Gigasporaceae, Glomeraceae and Paraglomeraceae, Table S1). OTU accumulation curves for both top- and subsoil reached a plateau, indicating that we captured the majority of the diversity (Fig. S2). We observed a highly significant community shift when comparing the top- and the subsoil (PERMANOVA, *F*_1,16_= 8.67, *P*<0.001, Fig. 3). Most remarkably, Claroideoglomeraceae and Diversisporaceae exhibit inversely proportional relative abundances across the studied soil profiles (Fig. 1). In topsoil OTUs assigned to Diversisporaceae represented 41.8% of the reads whereas in subsoil they represented 7.3% (GLM, *F*_1,15_=50.83 *P*<0.001). Conversely, OTUs assigned to Claroideoglomeraceae represented 15.0% of the reads in topsoil but 59.9% in subsoil (GLM, *F*_1,15_=17.87 *P*<0.001). The greater relative abundance of the family Claroideoglomeraceae in subsoil is not correlated with its nominal diversity but with a modest increase in relative richness (Fig. S3a,b). This finding may point to some species in the Claroideoglomeraceae family being subsoil specialists and particularly dominant in this compartment. This hypothesis is supported by previous results where Claroideoglomus etunicatum spores were more commonly found in deeper soil layers (Oehl et al., 2005; Yang et al., 2010). Conversely, both nominal and relative diversity in the Diversisporaceae decreased in subsoil showing no evidence for specialization in subsoil within this family. The family Glomeraceae however, is detected at a mostly constant relative abundance across topsoil (34.1%) and subsoil (28.4%), but decreases in abundance from rhizosphere (49.0%) to bulk soil and drilosphere considered together (23.6%; GLM, *F*_1,15_=6.79 *P*=0.02) (Fig. 1). Members of the Glomeraceae are known to preferentially allocate biomass inside the root while producing limited biomass in the soil (Powell et al., 2009). Our observations broadly support the idea that, due to producing a limited extraradical mycelium, Glomeraceae species are expected to preferentially colonize the direct surroundings of the root and to rapidly decrease in abundance outside the rhizosphere. We hypothesize that species with a preferentially intraradical lifestyle are less responsive to abiotic factors outside the root and therefore can readily colonize different soil horizons. In turn, those intraradical lifestyles would be mostly affected by host characteristics.

**Fig. 1.**
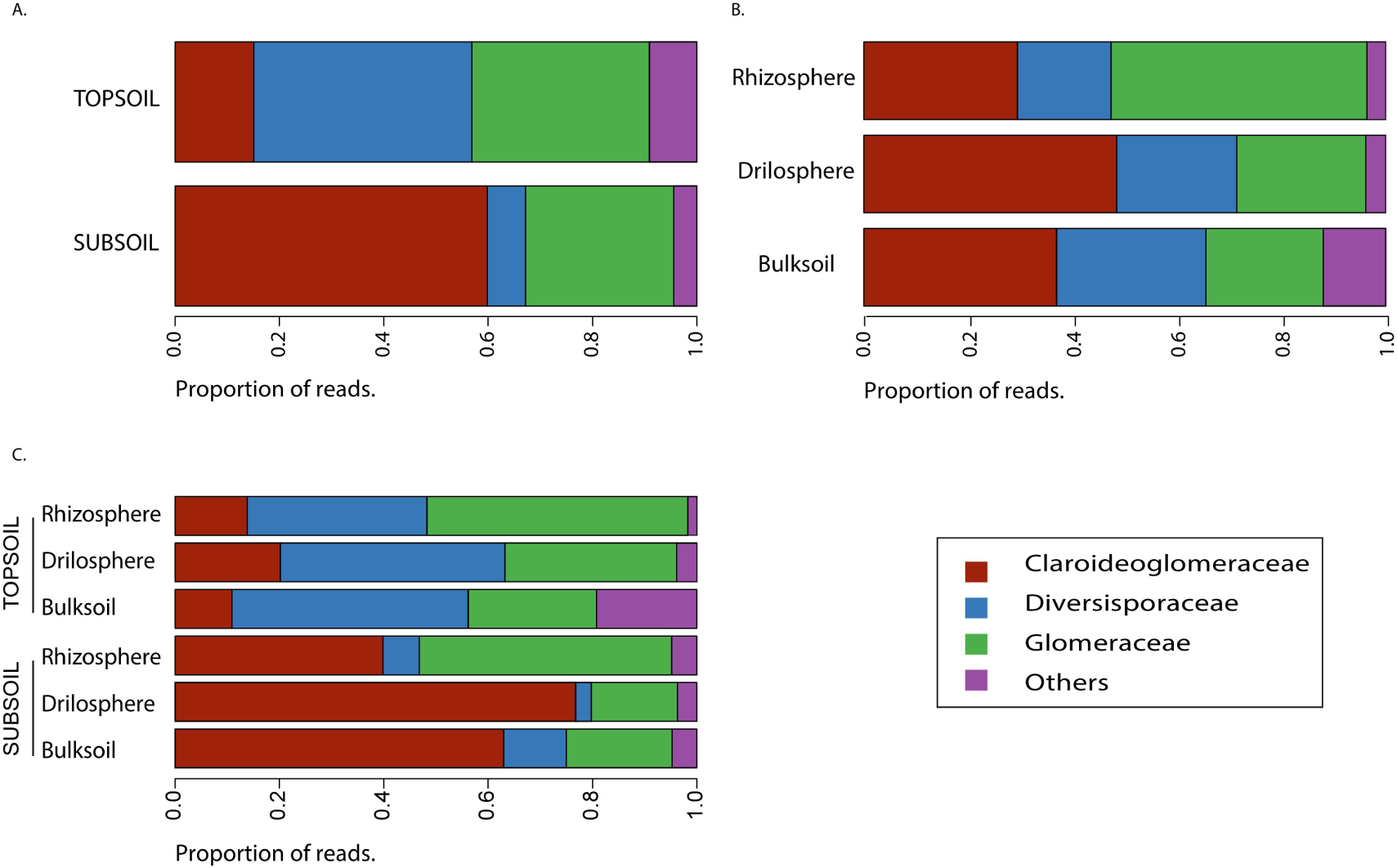
Relative abundance of reads per family in different soil compartments. Proportion of reads assigned to each family for depth (panel A), compartment (B) and depth and compartment (C). Families are coded by color. The category “Others” comprises the families Ambisporaceae, Archaeosporaceae, Gigasporaceae and Paraglomeraceae, as well as OTUs assigned only at the subphylum level. Topsoil = 10–30 cm, subsoil = 60–75 cm. Extended results are presented in the supplementary materia

**Fig. 2.**
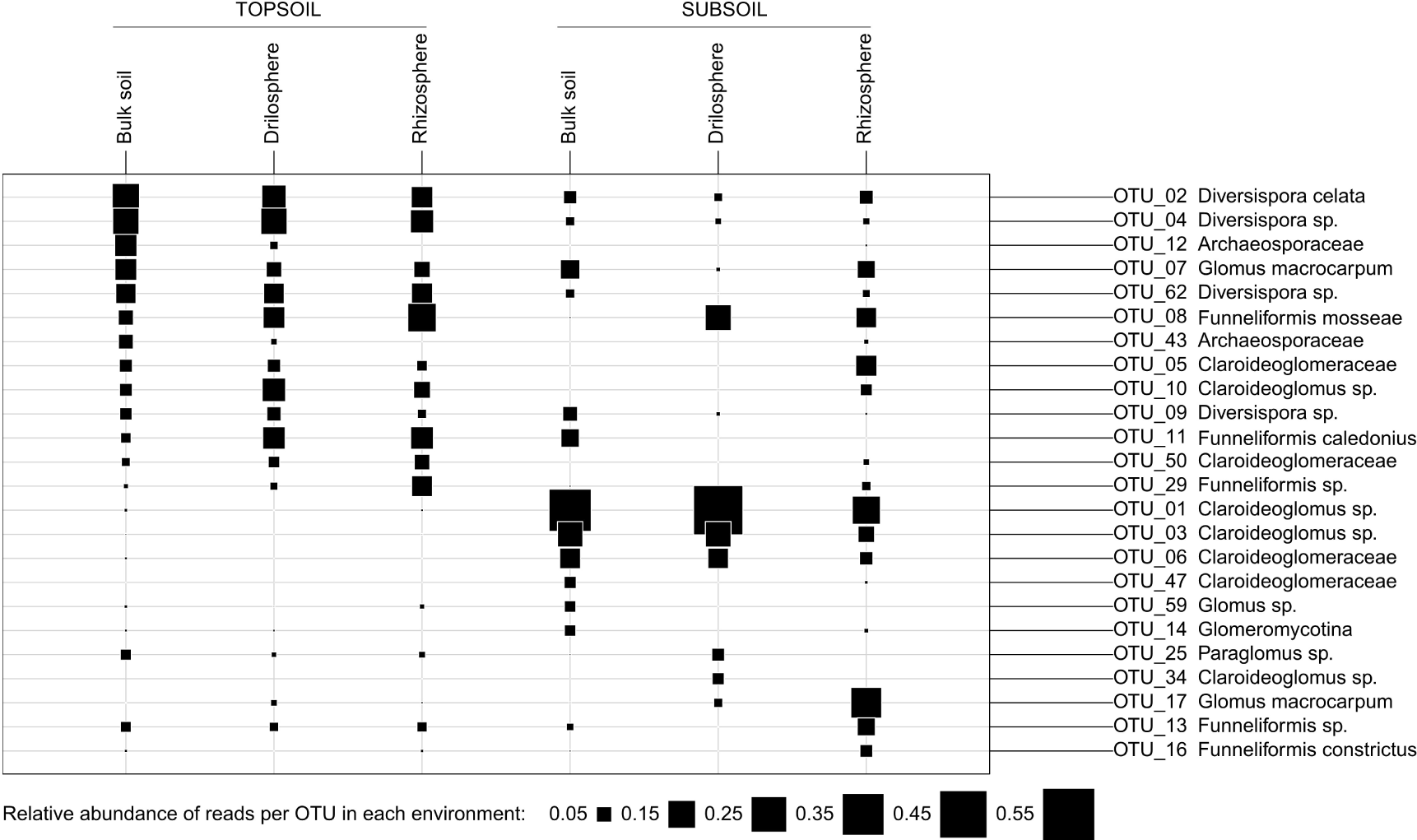
Relative abundance of the 10 most abundant OTUs for each soil compartment. Relative abundance of the OTU is represented by the area of the square (see scale below figure panel). The OTU list corresponds to the 10 most abundant OTUs for each environment. Topsoil = 10–30 cm, subsoil = 60–75 cm. Taxonomic assignment of each OTU is given (also see Table S1).

**Fig. 3.**
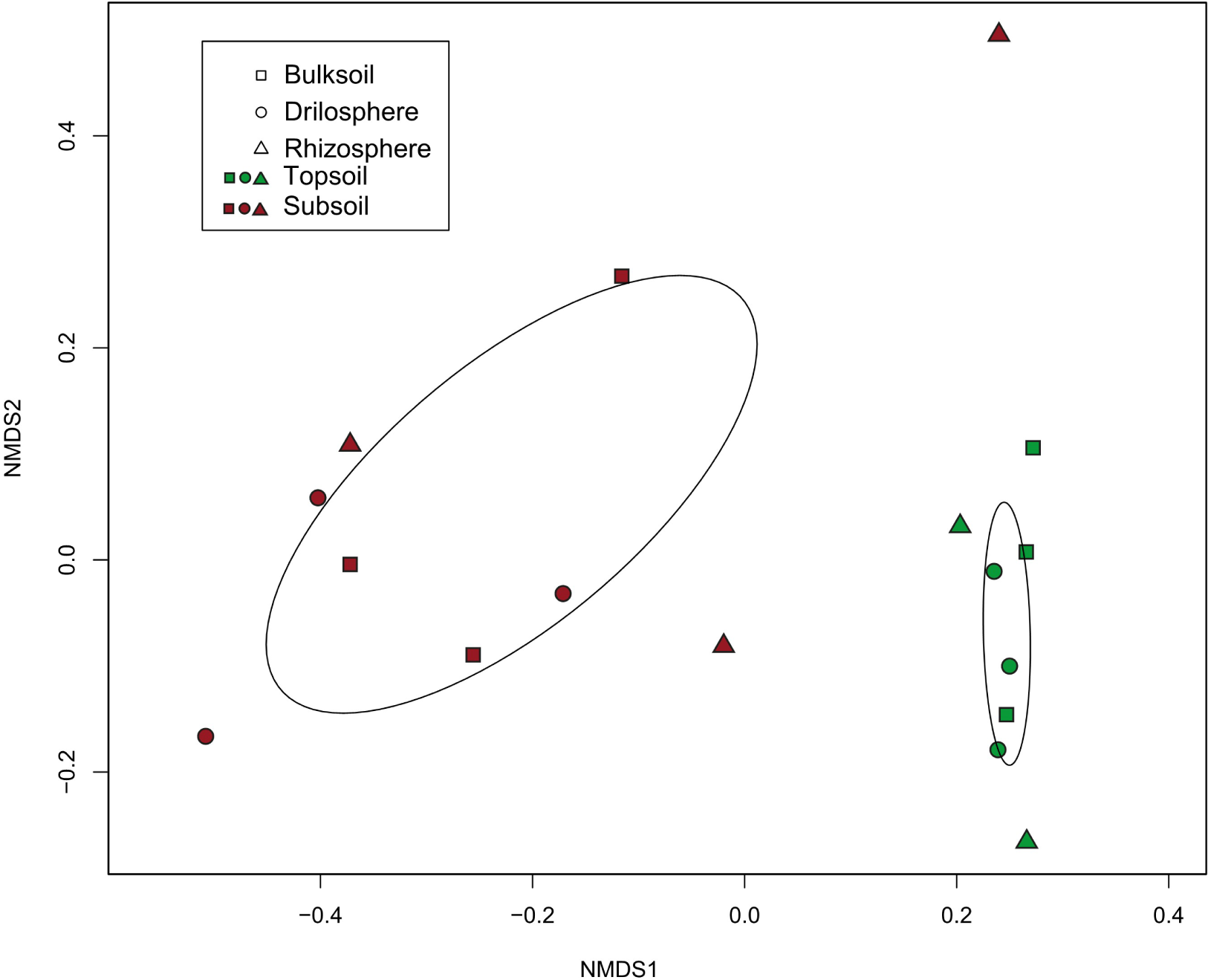
Community ordination of AMF in different soil compartments. Non-metric multidimensional scaling (NMDS) of Bray-Curtis pairwise community dissimilarities. The OTU table was normalized to the minimum amount of reads per sample. Ellipses represent one standard deviation around the centroid of each soil depth. Lines link each sample to the centroid of the group. Topsoil = 10–30 cm, subsoil = 60–75 cm. Depth is coded by color and compartment by symbol shape.

In our subsoil samples OTU richness (27.4 ± 5.9) was significantly lower than in topsoil (41.6 ± 6.0; GLM, *F*_1,15_=23.83 *P*<0.001). Nonetheless we detected a total of 49 OTUs in subsoil, with two OTUs (OTU_40 subphylum Glomeromycotina and OTU_68 genus Glomus) exclusively found in the subsoil, also before normalization (data not shown). Agricultural soils are subjected to a set of disturbances including fertilization or plant removal during harvest; topsoils are additionally subjected to high disturbance in form of tillage, negatively influencing AMF diversity (Kabir, 2005). Applying the C-S-R (competitor, stress tolerator, ruderal) framework to AMF (Chagnon et al., 2013), we would expect topsoil to be dominated by more ruderal species (i.e. elevated growth rates, rapid and abundant spore production, etc.) and subsoil by stress tolerators (i.e. low growth rates, long lived mycelium, etc.). We believe this may be one of the major factors explaining the observed differences in the communities across depth.

In our study no compartment effect was detected for the communities (PERMANOVA, F_2,16_= 0.68, P=0.66), besides the mentioned change in relative abundance of the family Glomeraceae (see above). However, previous studies conducted with the same samples show that bacterial communities exhibit a clear compartmentation in the subsoil (Uksa et al., 2015). This difference might be related to the linear, hyphal growth habit of fungi and their unique ability to integrate over larger soil volumes.

We were able to show that subsoil communities are clearly different and not only a subset of topsoil communities, and found contrasting patterns of abundance for different families. Whether this shift in community composition also means a shift in function remains unknown; but the clear difference in dominant families with depth suggests turnover also in functional traits (Powell et al. 2009). Our results emphasize the need to account for subsoil when designing agricultural management strategies and highlight the potential role of deeper soil layers as a biodiversity reservoir.

## Acknowledgements

MR acknowledges funding through the Federal Ministry of Education and Research (BMBF) initiative ‘BonaRes - Soil as a sustainable resource for the bioeconomy’ for the project Soil^3^. TC was supported by the project SENSE (Structure and Ecological Niche in the Soil Environment; EC FP7 - 631399 - SENSE). Illumina sequencing was carried out at the Berlin Center for Genomics in Biodiversity Research (BeGenDiv, Berlin, Germany). All authors thank Doreen Fischer and Marta Fogt for their involvement in the sampling and sample preparation processes, respectively. MS-H thanks Margarita Moreno Bayón for constant support.

